# Optimizing the PBS1 Decoy System to Confer Resistance to Potyvirus Infection in Arabidopsis and Soybean

**DOI:** 10.1101/2020.01.12.903575

**Authors:** Sarah E. Pottinger, Aurelie Bak, Alexandra Margets, Matthew Helm, Lucas Tang, Clare Casteel, Roger W. Innes

**Author notes:** **Corresponding author:** Roger W. Innes.

## Abstract

The *Arabidopsis* resistance protein RPS5 is activated by proteolytic cleavage of the protein kinase PBS1 by the *Pseudomonas syringae* effector protease AvrPphB. We have previously shown that replacing seven amino acids at the cleavage site of PBS1 with a motif cleaved by the NIa protease of turnip mosaic virus (TuMV) enables RPS5 activation upon TuMV infection. However, this engineered resistance conferred a trailing necrosis phenotype indicative of a cell death response too slow to contain the virus. We theorized this could result from a positional mismatch within the cell between PBS1^TuMV^, RPS5 and the NIa protease. To test this, we re-localized PBS1^TuMV^ and RPS5 to cellular sites of NIa accumulation. These experiments revealed that relocation of RPS5 away from the plasma membrane compromised RPS5-dependent cell death in *N. benthamiana*, even though PBS1 was efficiently cleaved. As an alternative approach, we tested whether overexpression of plasma membrane-localized PBS1^TuMV^ would enhance RPS5 activation by TuMV. Significantly, over-expressing the PBS1^TuMV^ decoy protein conferred complete resistance to TuMV when delivered by either *Agrobacterium* or by aphid transmission, showing that RPS5-mediated defense responses are effective against bacterial and viral pathogens. Lastly, we have now extended this PBS1 decoy approach to soybean by modifying a soybean PBS1 ortholog to be cleaved by the NIa protease of soybean mosaic virus (SMV). Transgenic overexpression of this soybean PBS1 decoy conferred immunity to SMV, demonstrating that we can use endogenous PBS1 proteins in crop plants to engineer economically relevant disease resistant traits.

## INTRODUCTION

The *Arabidopsis* disease resistance protein RESISTANCE TO PSEUDOMONAS SYRINGAE 5 (RPS5) confers resistance to *Pseudomonas syringae* through the detection of the effector protein AvrPphB (Simonich and Innes 1995). AvrPphB is a cysteine protease that targets serine-threonine kinases involved in pathogen-associated molecular pattern (PAMP)-triggered immunity (Zhang et al. 2010; Shao et al. 2003). One of these kinases, PBS1, forms a preactivation complex with RPS5 and triggers RPS5 activation upon AvrPphB-dependent cleavage (Ade et al. 2007). It is the resulting conformational change in PBS1 that activates RPS5, rather than the cleavage itself, as shown by the observation that insertion of three alanine residues into the cleavage loop of PBS1 is sufficient to activate RPS5 in the absence of cleavage (DeYoung et al. 2012). Therefore, RPS5 can theoretically confer resistance to any pathogen with an effector capable of causing the necessary conformational change in PBS1.

The work presented here follows on from that published in Kim et al. 2016 in which novel resistance to Turnip mosaic virus (TuMV) was engineered by insertion of a cleavage motif for the NIa protease of TuMV into PBS1 (PBS1^TuMV^). However, the engineered resistance was not complete, as transgenic PBS1^TuMV^ Arabidopsis plants displayed a trailing necrosis phenotype. This suggests the immune response produced by this engineered resistance was too slow to prevent the spread of viral infection (Kim et al. 2016). We theorized this could be due to a positional mismatch within the cell between the PBS1/RPS5 complex and the viral protease. Both PBS1 and RPS5 are tethered to the plasma membrane, along with AvrPphB (Qi et al. 2012; Dowen et al. 2009). However, the TuMV NIa protease is primarily located at the endoplasmic reticulum and in the nucleus (Cotton et al. 2009; Restrepo et al. 1990), raising the possibility that PBS1^TuMV^ is only cleaved late in the infection process, once high levels of viral proteins have accumulated and neighboring cells have already been infected. To test this hypothesis, we used genetic techniques to relocate the PBS1^TuMV^-RPS5 complex to regions of NIa accumulation and then assessed for improved activation of RPS5. We also tested whether overexpression of plasma membrane-localized PBS1^TuMV^ could enhance resistance. These experiments revealed that whilst the localization of PBS1^TuMV^ does not affect its ability to be cleaved by NIa protease, RPS5 is unable to induce cell death, unless it is located at the plasma membrane. However, overexpression of plasma membrane localized PBS1^TuMV^ conferred complete resistance to TuMV systemic infection, eliminating the trailing necrosis phenotype. We also show that the PBS1 decoy system can be deployed in crop plants, as transgenic overexpression of a soybean PBS1 protein engineered to be cleavable by the NIa protease of soybean mosaic virus (SMV) conferred immunity to SMV in soybean. Because the *PBS1* gene is highly conserved across angiosperms (Caldwell and Michelmore, 2009), this last finding has broad implications for engineering novel disease resistance traits in crops.

## RESULTS

### PBS1^TuMV^ and RPS5 can be relocated to various sub-cellular compartments by replacing their N-terminal acylation motifs with other sub-cellular targeting sequences

To test whether the PBS1/RPS5 system remains functional when not at the plasma membrane, we modified acylation motifs present in the N-termini of RPS5 and PBS1, which anchor these proteins to the plasma membrane (PM) (Qi et al. 2012 and 2014). Specifically, we generated dexamethasone-inducible constructs that targeted RPS5 and PBS1 to the cytoplasm, nucleus and to ER-derived membrane complexes induced under TuMV infection (see Methods). To ensure that any phenotypes observed were not due to the N-terminal location of the targeting sequence a C-terminal RPS5-NLS construct was also made. These constructs were then fused to super YFP and transiently expressed in *Nicotiana benthamiana*.

As shown in Figure 1, confocal microscopy analyses revealed that all re-localization constructs accumulated at subcellular sites other than the plasma membrane. Constructs containing a nuclear localization signal (NLS) produced a strong accumulation in the nucleus 24h after the induction of gene expression which the exception of RPS5-NLS, whose localization remained more cytoplasmic. The cytoplasmic (Cyt.) PBS1^TuMV^-sYFP construct produced a strong cytoplasmic accumulation as well as accumulation in the nucleus. However, cytoplasmic RPS5-sYFP accumulated to much lower levels (Fig. 1), consistent with the reduced stability of the RPS5^G2/3A,C4A^–sYFP construct that has been published previously (Qi et al. 2012).

**Fig. 1.**
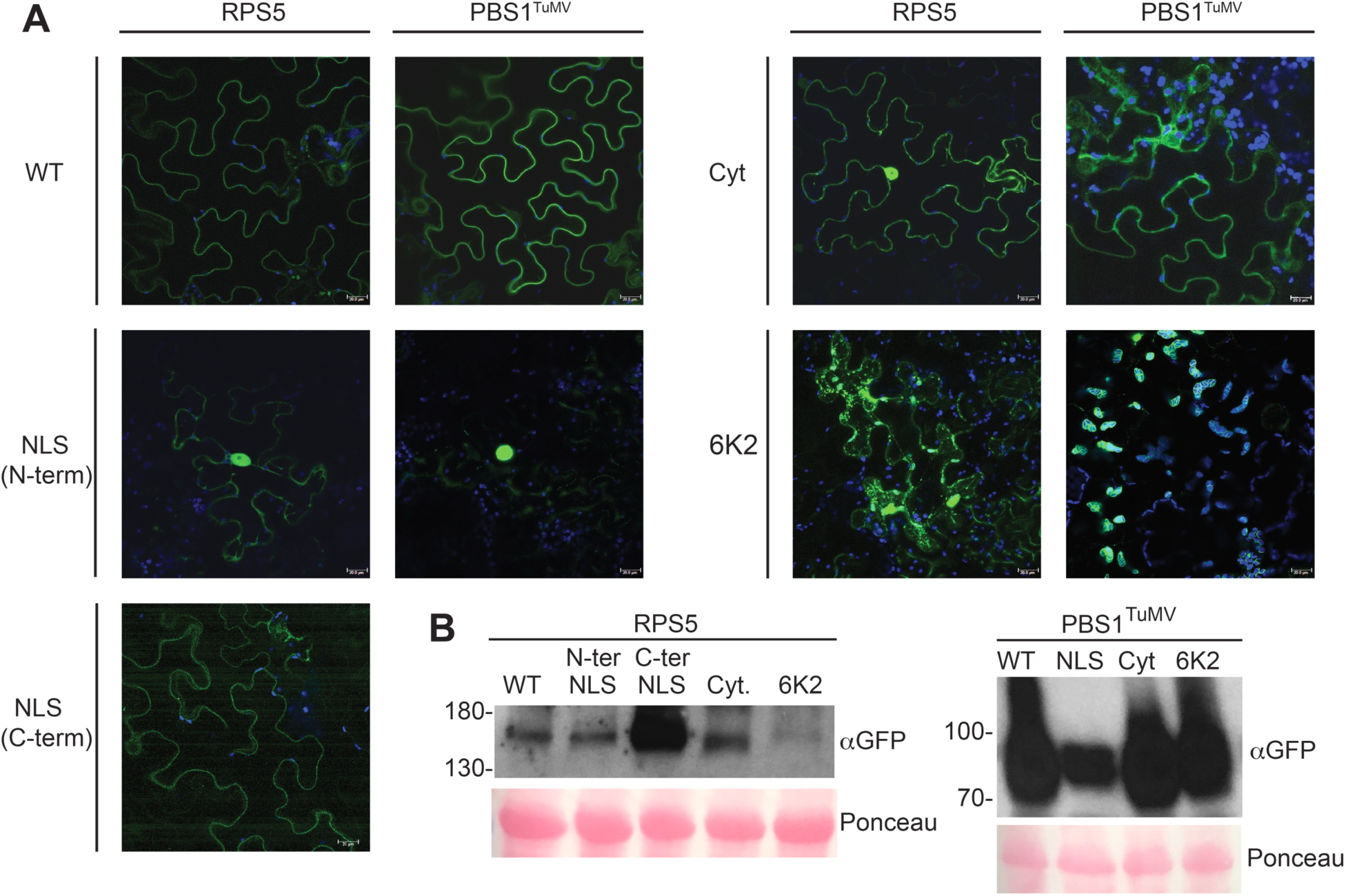
Subcellular localization of the PBS1^TuMV^ and RPS5 constructs. **A,** Fluorescence images showing the expression and localization of re-localized PBS1^TuMV^-sYFP and RPS5-sYFP at 24 hours after the induction of gene expression in *N. benthamiana*. Each image is a Z-stack comprised of 5 single sections. The scale bar represents 20µm. Each construct was imaged independently three times. **B,** Immunoblot showing protein accumulation at 24 hours post induction of gene expression.

To more fully match the localization of the PBS1/RPS5 complex to sites of viral replication, the acylation sites of PBS1^TuMV^ and RPS5 were replaced with the complete TuMV 6K2 protein sequence. Under conditions of viral infection, the 6K2 protein has been shown to manipulate host endomembranes into viral ‘replication factories’ (Cotton et al. 2009). We predicted that fusion of the 6K2 protein to the N-termini of PBS1^TuMV^ and RPS5 would localize them to these viral replication factories during infection, and possibly to ER-derived structures in the absence of virus. The 6K2-PBS1^TuMV^-sYFP accumulated on the surface of chloroplasts and caused chloroplasts to clump (Fig. 1 and Supplementary Fig. S1) as has been previously shown for 6K2-YFP (Wei et al. 2010). 6K2-RPS5-sYFP accumulated in vesicle-like puncta within the cytoplasm (Fig. 1). Together these data suggest that RPS5 and PBS1^TuMV^ can be re-localized to other subcellular compartments.

### Re-localized PBS1^TuMV^ is still cleaved by the TuMV NIa protease

As we have shown that PBS1^TuMV^ can be re-localized, we next wanted to test if PBS1^TuMV^ is still capable of interacting with, and being cleaved by, the TuMV NIa protease in its new location. All re-localized PBS1^TuMV^ constructs showed a robust expression at 6h after dexamethasone induction; however, the NLS-PBS1^TuMV^ construct accumulated to a lower level (Fig. 2). We therefore assessed whether these PBS1^TuMV^ derivatives could be efficiently cleaved by TuMV NIa protease in their new locations. As previously shown, PM-localized PBS1^TuMV^ was readily cleaved by the TuMV NIa protease at 6h post induction (Fig. 2; Kim et al. 2016). NLS-PBS1^TuMV^ was also cleaved, but its cleavage products accumulated to lower levels compared to that of PM-localized PBS1^TuMV^ cleavage products. These data suggest that PBS1^TuMV^-HA cleavage products may be more rapidly degraded in the nucleus. Both the 6K2- and cytoplasmic localized PBS1^TuMV^-HA cleavage products displayed accumulation similar to wild type. The observation that PBS1^TuMV^ could be cleaved regardless of sub-cellular location indicates that TuMV NIa is present in all of these locations, at least when overexpressed in *N. benthamiana*.

**Fig. 2.**
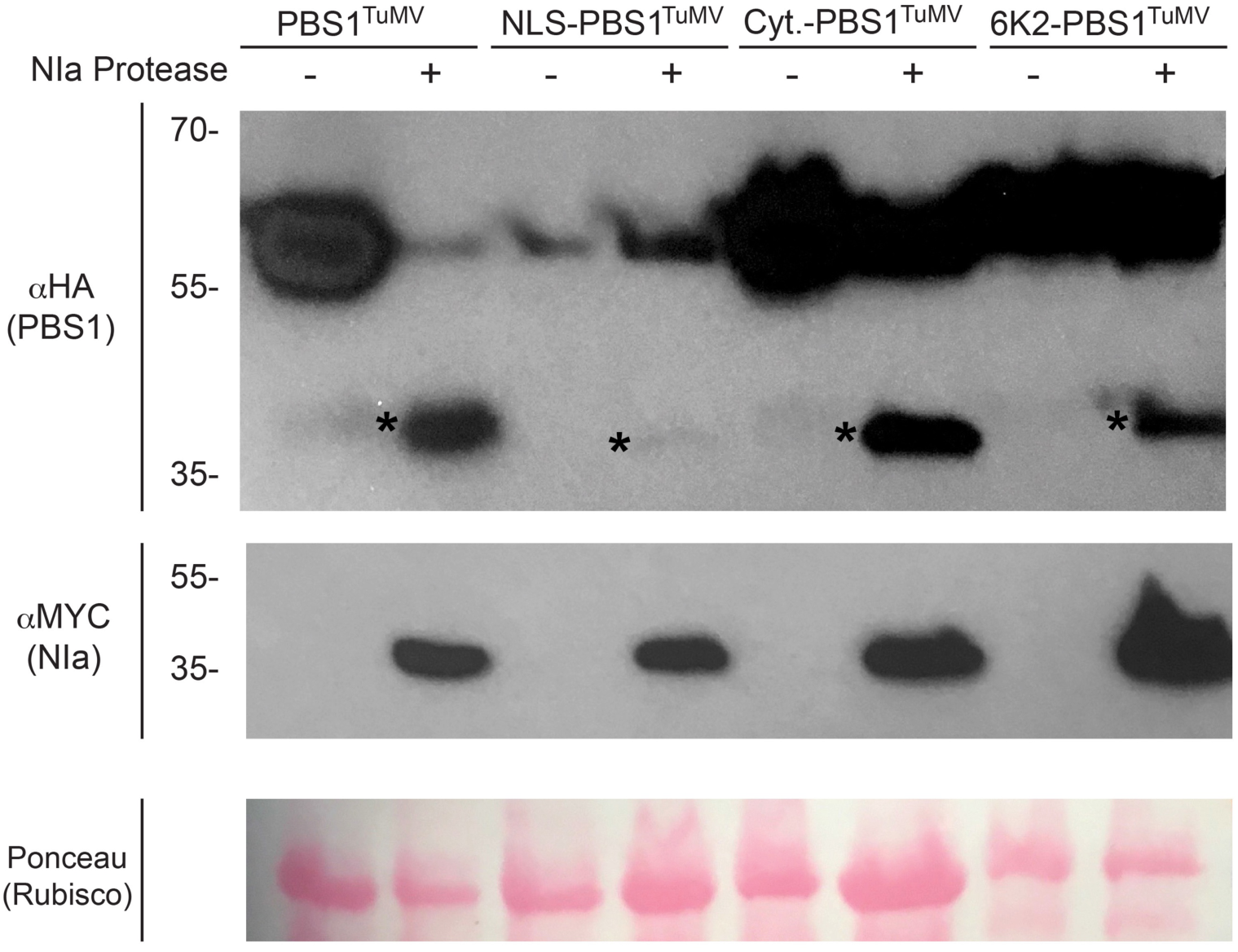
TuMV NIa-mediated cleavage of the re-localized PBS1^TuMV^ constructs in *N. benthamiana*. Immunoblots showing the expression of re-localized PBS1^TuMV^-HA constructs and their cleavage in the presence of MYC-tagged Turnip Mosaic Virus NIa Protease. At six hours after the induction of gene expression total protein was extracted and immunoblotted. Ponceau S-stained blots are shown to assess variability in sample loading. * indicates bands corresponding to PBS1 cleavage products.

### Re-localized RPS5/PBS1^TuMV^ complexes are unable to activate programmed cell death

Having confirmed that re-localized PBS1^TuMV^-HA is efficiently cleaved by the TuMV NIa protease, we next tested whether co-localized PBS1^TuMV^ and RPS5 could still activate programmed cell death when transiently co-expressed in *N. benthamiana*. The PM-localized RPS5/PBS1^TuMV^ complex induced a robust cell death in *N. benthamiana* leaves when co-expressed with TuMV NIa protease (Fig. 3A). However, none of the re-localized RPS5/PBS1^TuMV^ complexes were able to do so (Fig. 3A), suggesting that RPS5 may not function in these locations. Immunoblot analyses revealed that the relocalized RPS5 constructs accumulated to levels similar to that of PM-localized RPS5, with the exception of 6K2-RPS5, which was lower (Fig. 3B). Thus, the lack of cell death induction was unlikely due to insufficient levels of RPS5.

**Fig. 3.**
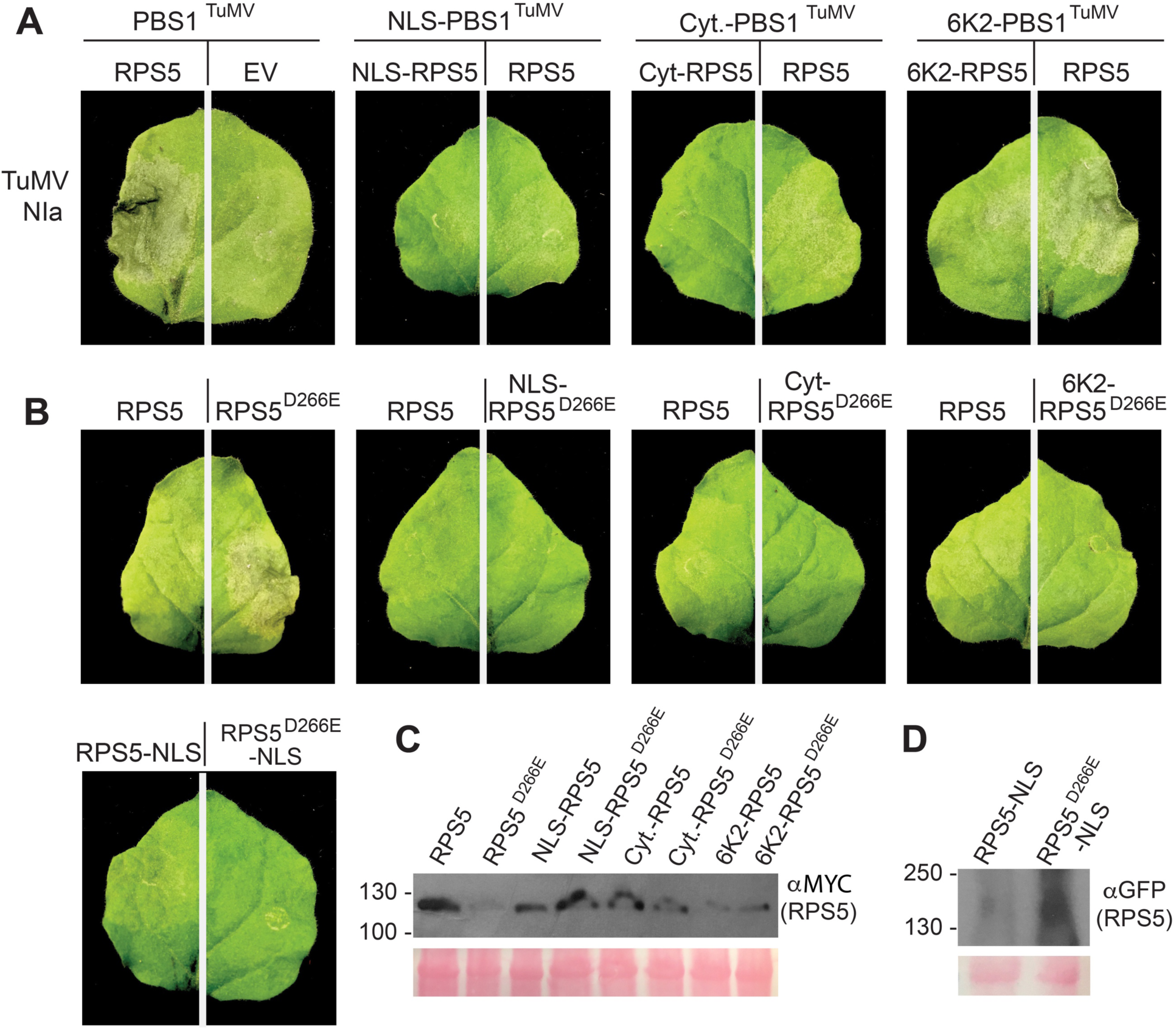
Re-localized RPS5 is unable to signal for cell death. **A and B,** Photographs showing leaf phenotypes elicited in response to TuMV NIa protease by PM and re-localized PBS1^TuMV^ and RPS5 (A), as well as re-localized auto-active RPS5 (RPS5^D266E^) (B), at 24 hours after the induction of gene expression in transient transformed *N. benthamiana* leaves. The leaves shown are representative of the phenotype seen in all 15 inoculated leaves. **C**, Immunoblot showing the protein accumulation of WT and D266E RPS5-Myc constructs (used for the inoculations shown in the top two rows) when expressed by themselves. Accumulation of PBS1^TuMV^-HA and TuMV NIa-Myc constructs used in this assay can be seen in Figure 2. **C,** Immunoblot showing the protein accumulation of WT and D266E YFP-RPS5-NLS constructs used in the bottom left images.

To confirm that the lack of cell death was due to an inability of re-localized RPS5 to signal from locations other than the PM, we used an auto-active mutant of RPS5 containing a D266E substitution (RPS5^D266E^), which constitutively activates cell death in the absence of PBS1 (Ade et al. 2007). Significantly, re-localized RPS5^D266E^ was unable to induce cell death when transiently expressed in *N. benthamiana,* despite accumulating to higher levels than PM-localized RPS5^D266E^, while PM-localized RPS5^D266E^ triggered a strong cell death response (Fig. 3). This suggests that the failure of the re-localized PBS1^TuMV^/ RPS5 complex to induce cell death is likely a result of RPS5 requiring PM localization to activate cell death, rather than an inability of cleaved PBS1^TuMV^-HA to activate RPS5.

The observation that RPS5 must reside at the PM to induce cell death lead us to test whether re-localized PBS1^TuMV^ can activate PM-localized RPS5, which may enable a more rapid detection of TuMV NIa protease than previously seen in the PBS1 decoy system (Kim et al. 2016) and thus overcome the trailing necrosis phenotype. We thus tested each modified PBS1^TuMV^ construct for activation of wild-type RPS5; however, only the 6K2-PBS1^TuMV^ construct showed a cell death response (Fig. 3A). These results indicate that re-localization of PBS1 away from the PM eliminates its ability to activate RPS5-dependent HR, with the exception of the 6K2-PBS1^TuMV^ construct. Notably, this is the construct that localized around the periphery of chloroplasts (Fig. 1 and Supplementary Fig. S1), and perhaps reflects the close association of chloroplasts with the PM.

### Expression of 6K2-PBS1^TuMV^ in *Arabidopsis* does not complement a *pbs1* loss of function mutation

As 6K2-PBS1^TuMV^ could induce cell death in *N. benthamiana* when co-expressed with WT RPS5 and TuMV NIa protease, we proceeded to assess whether this construct could confer resistance to TuMV infection in transgenic *Arabidopsis* plants. Transgenic lines expressing 6K2-PBS1^TuMV^ under a 35S promoter were generated in *pbs1* and *rps5* mutant backgrounds. The *rps5* background was used to assess whether any observed resistance was dependent on RPS5, whilst the *pbs1* line was used to eliminate competition with the WT PBS1 for RPS5 binding. When these lines were infected with TuMV (6K2-GFP), systemic spread of TuMV was observed three weeks after inoculation (Fig. 4A). Systemic infection was verified by immunoblot analysis (Fig. 4B). This result, combined with the transient expression assays described above, indicate that re-localization of PBS1^TuMV^ to sites of viral replication does not enhance the efficacy of the RPS5-PBS1 decoy system.

**Fig. 4.**
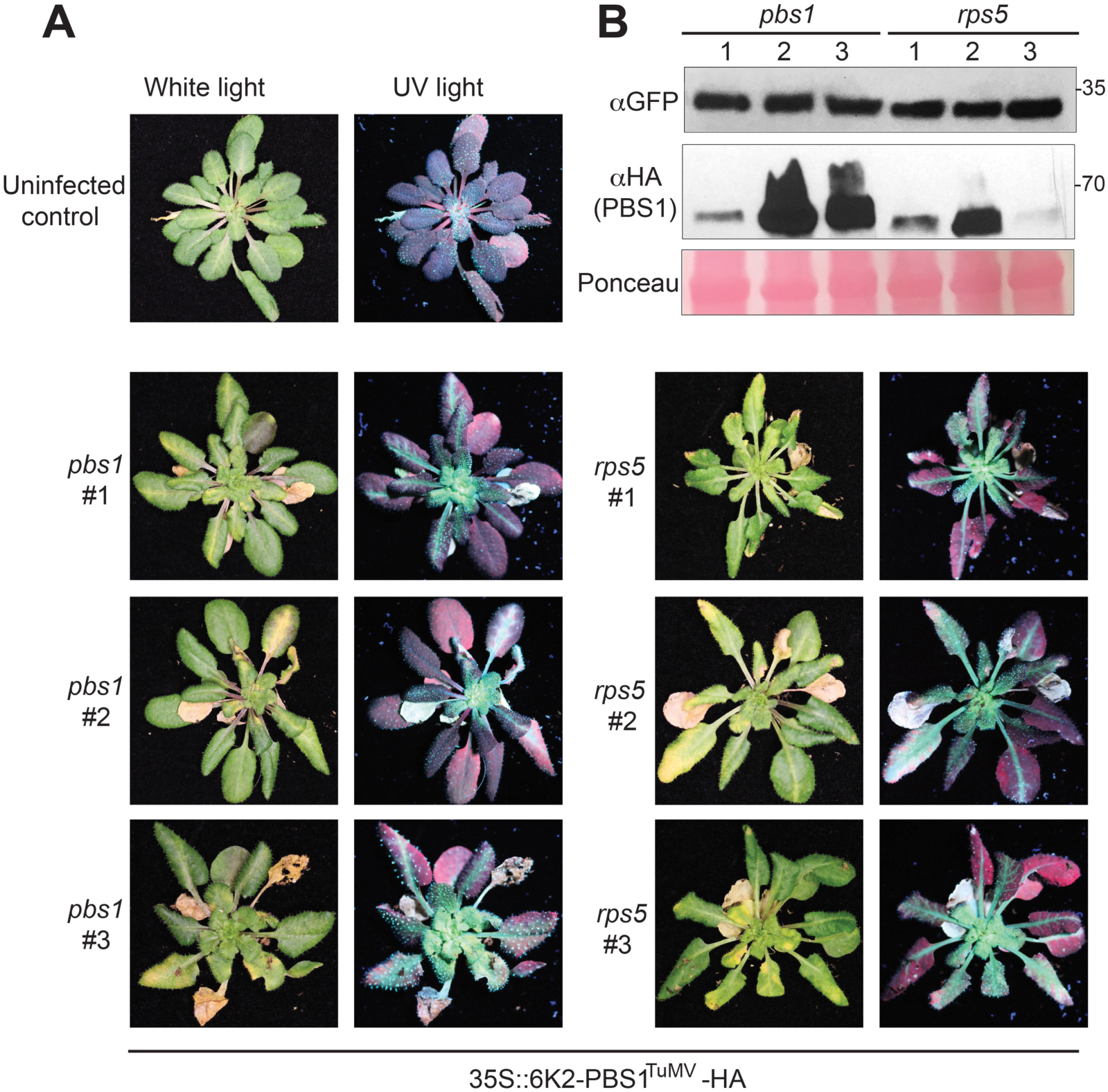
6K2-PBS1^TuMV^ does not function in transgenic *Arabidopsis*. **A,** White light and UV photographs showing the spread of TuMV(6K2-GFP) in three independent 6K2-PBS1^TuMV^ transformed homozygous lines in *pbs1* and *rps5* mutant backgrounds at three-weeks post infection. **B,** Immunoblot showing 6K2-PBS1^TuMV^ and virally produced 6K2-GFP expression levels in the lines pictured in A.

### Overexpression of PBS1^TuMV^ in *Arabidopsis* confers complete immunity to TuMV

As an alternative approach, we tested whether overexpressing PM-localized PBS1^TuMV^ in *Arabidopsis* would enhance resistance and ameliorate the trailing necrosis phenotype, as a correlation was observed previously between TuMV resistance and the level of PBS1^TuMV^ expression (Kim et al. 2016). Three independent homozygous lines expressing HA tagged PBS1^TuMV^ under a 35S promoter were generated in wild-type Col-0, *pbs1* and *rps5* backgrounds. As with the 6K2-PBS1^TuMV^ lines, the *rps5* background was used to confirm dependency on RPS5, and the *pbs1* line was used to ensure there would be no loss of signaling efficiency through competition with the WT PBS1 protein for association with RPS5. The stable overexpression of PBS1^TuMV^ in these lines appeared to have no fitness consequences, as plants grew similarly to wild type and, with the exception of the *pbs1* lines 1-5 and 2-1, produced an equivalent amount of seed (Supplementary Fig. S2).

Transgenic lines were inoculated with TuMV (6K2-GFP) using Agrobacterium-mediated delivery and assessed for TuMV accumulation in the non-inoculated, systemic leaves via ultra-violet light imaging at three weeks post infection (Fig. 5A). Consistent with our hypothesis, wild type Col-0 and *pbs1* genotypes over-expressing PBS1^TuMV^ displayed complete resistance to TuMV. In contrast, the *rps5* lines over-expressing PBS1^TuMV^ showed systemic GFP fluorescence derived from virally produced 6K2-GFP, indicative of successful TuMV infection (Fig. 5A). The lines expressing PBS1^TuMV^ under a native promoter showed the trailing necrosis phenotype previously published (Kim et al. 2016). These data establish that TuMV resistance in the wild type and *pbs1* transgenic lines is dependent on RPS5 expression and the expression level of PBS1^TuMV^. These results were verified using immunoblot analyses (Fig. 5B). We thus conclude that the trailing necrosis phenotype originally observed in the PBS1/RPS5 decoy system (Kim et al. 2016) can be overcome through overexpression of the decoy protein. Furthermore, expression of PBS1^TuMV^ in a Col-0 background does not compromise the recognition of AvrPphB. It can be seen in Fig. 6 that both the 35S and native promoter PBS1^TuMV^ lines in a Col-0 background were capable of AvrPphB recognition, whereas the lines in the *rps5* background showed a response to AvrPphB comparable to that of the empty vector. Interestingly, lines in the *pbs1* background showed a variable level of HR, suggesting that PBS1^TuMV^ may still be able to be cleaved by AvrPphB at a low level. These data indicate that the pool of RPS5 available for partnering with decoy PBS1 proteins is not severely limiting, thus it should be possible to expand the recognition specificity of RPS5 through the addition of multiple different decoy PBS1 proteins.

**Fig. 5.**
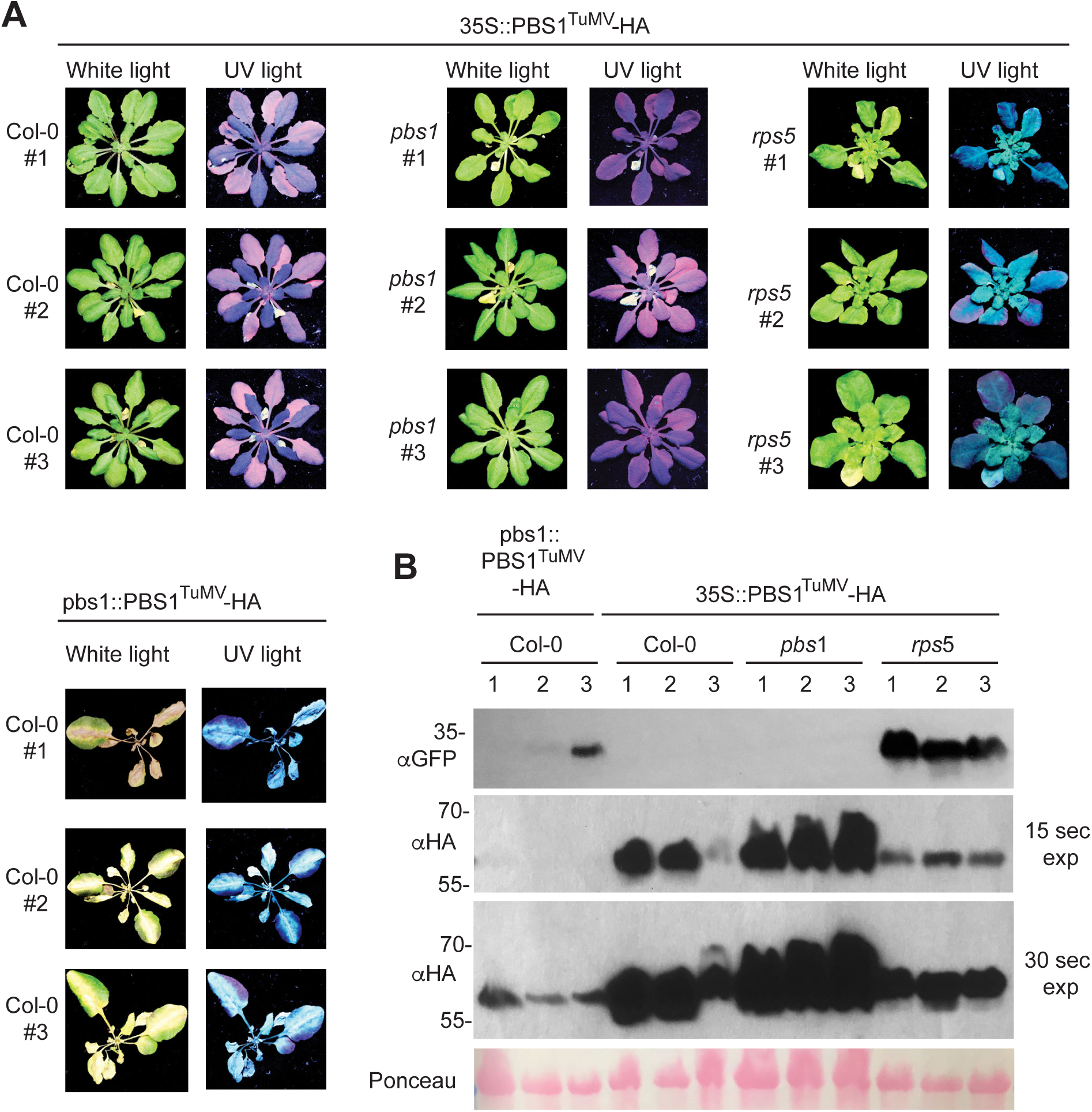
Overexpression of PBS1^TuMV^ confers complete immunity to TuMV. **A,** White light and UV photographs showing the spread of TuMV infection in homozygous 35S:PBS1^TuMV^ transformants in Col-0, *pbs1* and *rps5* backgrounds and pbs1::PBS1^TuMV^ in Col-0 at three weeks post infection by Agrobacterium-mediated delivery of viral RNAs. **B,** Immunoblot showing PBS1^TuMV^ and virally produced 6K2-GFP expression levels in the lines pictured in A.

**Fig. 6.**
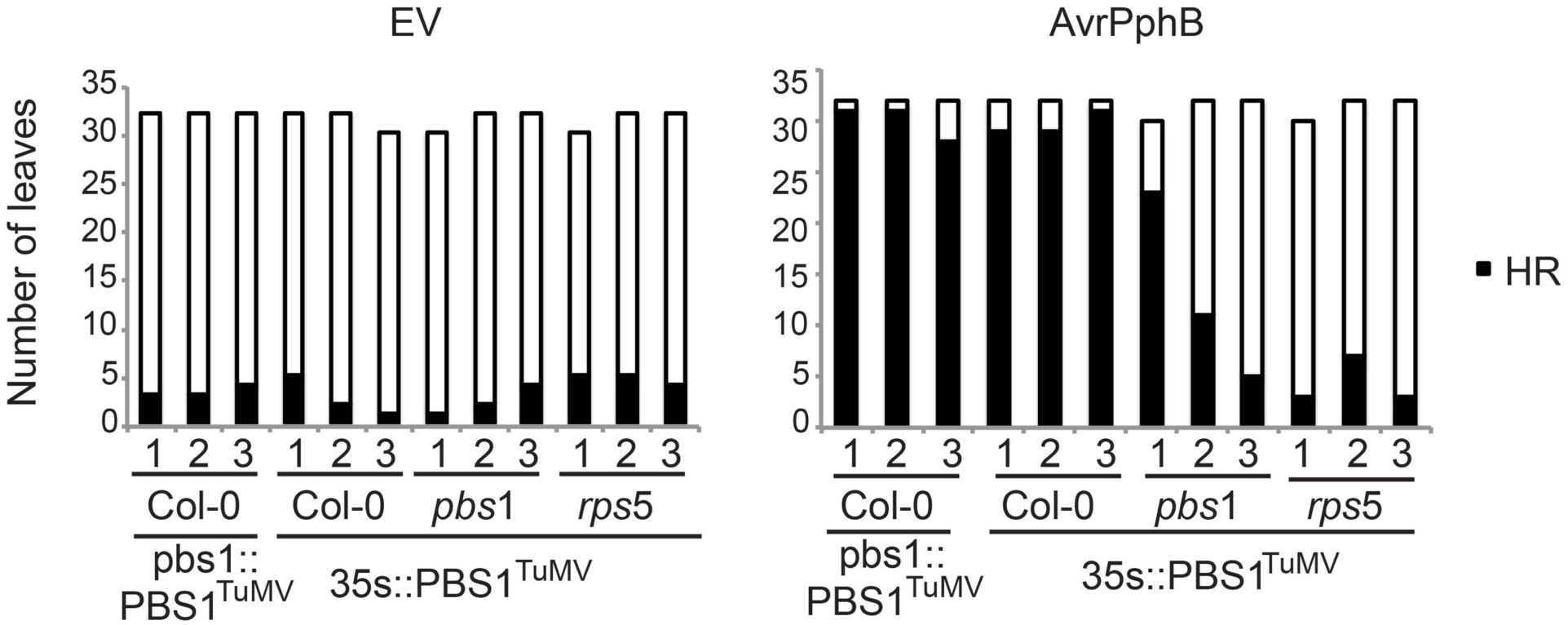
Arabidopsis expressing both WT PBS1 and PBS1^TuMV^ retain recognition of AvrPphB. Bar charts showing the number of leaves expressing an HR phenotype in response to infiltration with *Pseudomonas syringae* expressing either AvrPphB or an empty vector. Bar height represents the number of leaves infiltrated. The black bar represents the number of leaves expressing cell death. HR tests were done in four week old homozygous 35S:PBS1^TuMV^ transformants in Col-0, *pbs1* and *rps5* backgrounds and pbs1:PBS1^TuMV^ in Col-0.

To confirm that overexpression of PBS1^TuMV^ confers resistance to TuMV, we also conducted infection assays using aphid transmission, which is the natural vector for TuMV infection (Shattuck 2010). Homozygous lines expressing 35S:PBS1^TuMV^ were inoculated with aphids that had recently been feeding on TuMV (6K2:GFP)-infected plants (Walsh and Jenner 2002). Significantly, the *pbs1* lines over-expressing PBS1^TuMV^ displayed no observable viral symptoms two weeks post infection (Fig. 7A). In Col-0 lines, viral symptoms were only observed in the final of three replicates, in 2 of the 10 plants tested. Results from the final replicate are shown. The *rps5* lines, however, showed a strong GFP fluorescence caused by TuMV produced 6K2-GFP, as expected. These results were verified using immunoblot analysis (Fig. 7B). Here PBS1^TuMV^ expression levels are shown to be more variable, possibly due to increased protein turnover during aphid infection. However, these data still demonstrate that the PBS1^TuMV^ decoy is capable of conferring strong resistance to TuMV in a natural context.

**Fig. 7.**
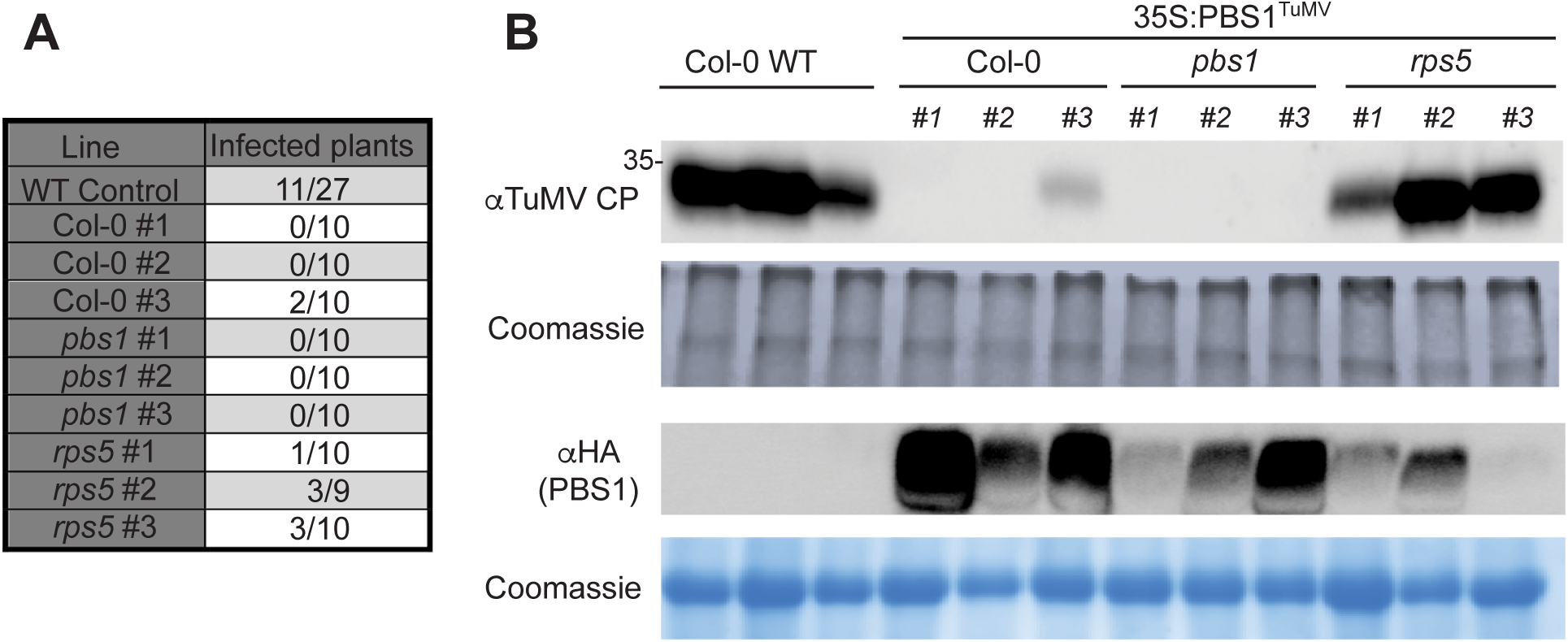
Aphid transmission of TuMV is blocked in PBS1^TuMV^ overexpressing lines. **A,** Table showing the percentage of *Arabidopsis* plants expressing GFP as a result of TuMV infection in Col-0, *pbs1* and *rps5* backgrounds at 15 days post aphid inoculation. **B,** Immunoblot showing PBS1^TuMV^ and TuMV Coat Protein expression levels in the lines represented in A.

### Overexpression of a soybean PBS1 decoy protein, *Gm*PBS1-1^SMV^, in soybean confers immunity to soybean mosaic virus

We have recently shown that PBS1 proteins from soybean can also be engineered to confer recognition of the NIa protease from SMV and that cleavage of these decoy proteins activates cell death in soybean protoplasts, suggesting the decoy engineering approach can be extended to a crop plant (Helm et al., 2019). It was unclear, however, whether transgenic soybean expressing the *Gm*PBS1-1^SMV^ decoy protein would confer resistance to SMV. We thus generated transgenic soybean (cv. Williams 82) constitutively expressing HA-tagged *Gm*PBS1-1^SMV^ under an enhanced 35S promoter. Three independent T0 soybean plants were obtained. Screening of T1 progeny from each line for expression of β-glucuronidase (see Methods), a marker on the T-DNA vector used for transformation, revealed that only one T1 family segregated the T-DNA in a 3:1 ratio. We harvested seed from three sibling T1 progeny of this line to generate three T2 families, two of which contained the T-DNA (families A and B) and one of which did not (family C), based on staining for GUS expression and genotyping by PCR. To test for resistance to SMV infection, unifoliate leaves of ∼12-day old T2 soybean plants were rub inoculated with SMV-Nv::GFP virions. Systemic spread of SMV was assessed by scoring the third trifoliate leaves for virus presence three weeks after inoculation. No observable mosaic symptoms or SMV coat protein (CP) accumulation was observed in families A or B (10 plants each), while 7 of 10 family C plants showed virus accumulation (Fig. 8). Consistent with these results, immunoblot analysis revealed that all plants in families A and B expressed PBS1-1^SMV^, while none of the family C plants showed PBS1^SMV^ expression (Fig. 8). These data indicate that transgenic overexpression of the *Gm*PBS1-1^SMV^ decoy protein confers resistance to systemic infection by SMV without a trailing necrosis phenotype, nor any obvious deleterious effects on plant growth.

**Fig. 8.**
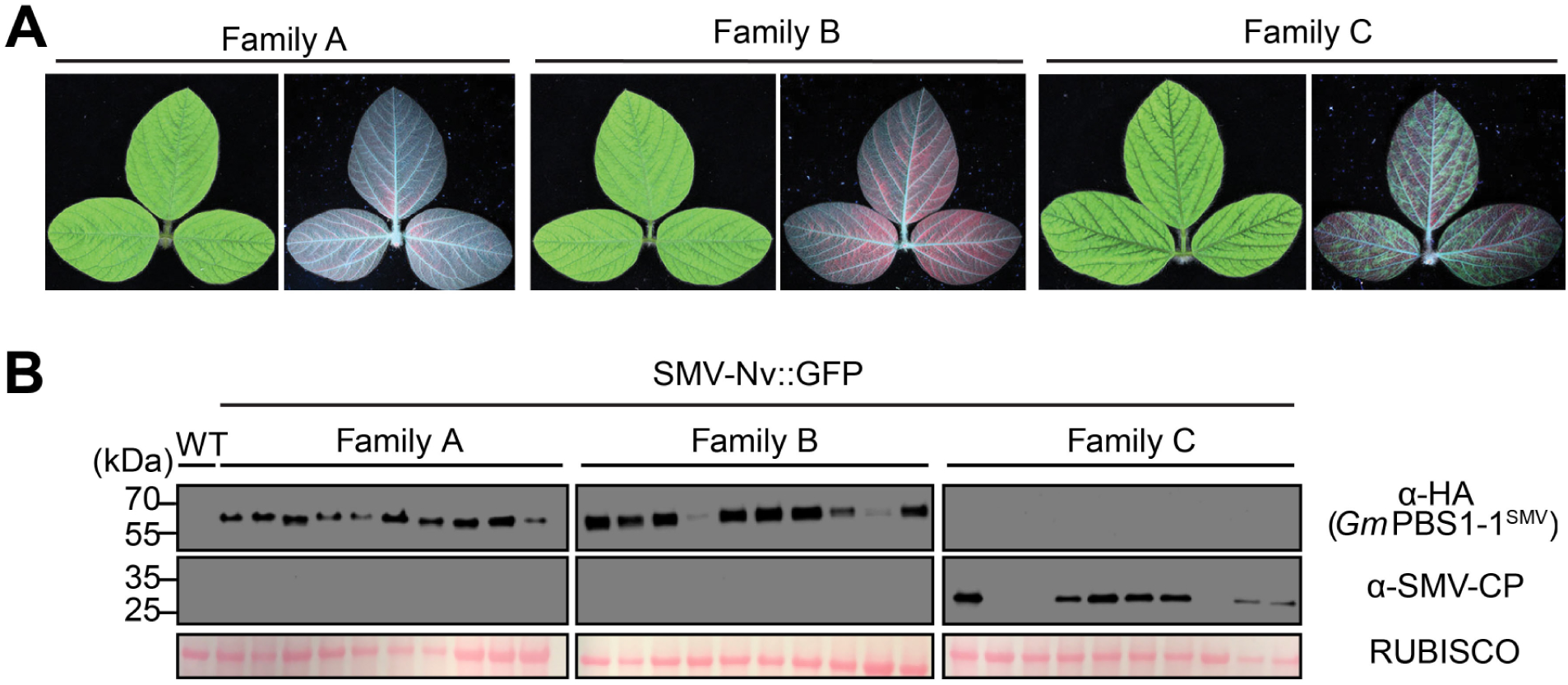
Overexpression of *Gm*PBS1-1^SMV^ in soybean confers resistance to SMV. **A,** White light and ultraviolet light photographs of soybean third trifoliate leaves from three sibling T3 families rub inoculated with SMV-Nv::GFP virions on their unifoliate leaves three weeks prior. Family C lacks the *PBS1-1^SMV^* transgene. White light images of adaxial sides (left) show SMV symptoms. Ultraviolet images of abaxial sides (right) show SMV accumulation as indicated by green fluorescence. Red color is from chlorophyll autofluorescence. **B,** Immunoblots of soybean trifoliate leaves from 10 individual plants from each family showing *Gm*PBS1-1^SMV^ expression (top), SMV coat protein (middle), and Ponceau stain of RUBISCO (bottom). Uninfected non-transgenic Williams82 (WT) was used for a negative control to show antibody specificity.

## DISCUSSION

The above experiments show that the PBS1 decoy system is capable of conferring complete resistance to a viral pathogen. We have also shown that plasma membrane localization is not essential for stable accumulation of PBS1^TuMV^, nor is it necessary for cleavage of PBS1^TuMV^ by the TuMV NIa protease. We have also shown that although cytosolic RPS5 is unstable, targeting to it to other cellular compartments can stabilize it. However, away from the plasma membrane, RPS5 cannot activate cell death, even when constitutively active.

There appears to be no general rule for the subcellular localization of NLR proteins other than their co-localization with their cognate effector. For example, the resistance protein L6 localizes to the Golgi and forced re-location to the vacuolar membrane did not impair its ability to function (Takemoto et al. 2012). It is known that disease resistance proteins such as MLA10 and Rx endogenously relocate from the cytosol to the nucleus, upon activation (Shen et al. 2007; Slootweg et al. 2010). However, while a nuclear localization enables MLA10 to interact with transcriptional activators and repressors to produce a defense response (Chang et al. 2013), Rx function in the nucleus is impaired (Slootweg et al. 2010). Furthermore, if the cognate effector of Rx is forced to accumulate in the host nucleus, Rx is unable to activate defense responses. This suggests that both Rx activation and signaling takes place in the cytoplasm, with nuclear accumulation possibly representing a form of negative regulation (Tameling et al. 2010). This nucleo-cytoplasmic partitioning is shared by RPS4. However, RPS4 activates distinct signaling pathways in the cytoplasm and the nucleus. Receptor recognition in the nucleus is capable of inhibiting bacterial growth, whilst being unable to activate the cell death response. However, recognition in the cytoplasm is capable of eliciting an HR while having minimal effects on bacterial growth (Heidrich et al. 2011). Both RPS5 and RPM1 are localized to the plasma membrane and do not relocate upon activation (Qi et al. 2012; Gao et al. 2011). However, there must be a signal that is translocated to the nucleus in order to alter gene expression and bring about the hypersensitive response, as it has been shown that compounds that disrupt transcription, like Amanitin, and translation, such as blasticidin and cyclohexamide, can inhibit HR. (Keen et al. 1981; He et al. 1993).

Our data indicate that RPS5 can only activate cell death pathways when located at the PM. Recent structural analyses of the Arabidopsis disease resistance protein ZAR1 provide functional insight into why this may be (Wang et al. 2019). ZAR1 oligomerizes upon activation and associates with the plasma membrane via its N-terminal coiled-coil domain. Oligomerized ZAR1 is then predicted to insert into the plasma membrane to form a pore that could act as a calcium channel. The resulting calcium influx would then lead to gene activation and cell death (Wang et al. 2019). It is known that calcium signaling is one of the first events triggered by effector recognition. Calcium channel blockers such as LaCl can inhibit the hypersensitive response induced by other PM-localized NLR proteins such as RPM1 (Grant et al. 2000). The nature of the calcium burst is also fine-tuned to the pathogen that elicits it, suggesting a close tie between calcium response and pathogen recognition (Grant et al. 2000). It is possible that RPS5 functions in a similar way, removing the need for downstream signalling partners by creating a calcium burst directly.

Alternatively, it may be that another protein is activated along with RPS5 and performs a signaling function. This putative partner of RPS5 would likely be located on the plasma membrane, as a cytoplasmic partner would be able to associate with RPS5 when localized to other cellular compartments. It is likely that RPS5 is capable of recruiting the signaling partner alone, i.e. without the aid of *At*PBS1, as it has been shown that autoactive RPS5 can cause HR when transiently expressed alone in *N. benthamiana* (Ade et al. 2007). It may be that the signaling partner and RPS5 act as a base for the formation of a higher-order signaling complex, as happens with the NLR proteins NLRC4 and NIAP2 in the activation of the mammalian inflammasome (Hu et al. 2015). In support of this model, RPS5 is known to form dimers or oligomers (Ade et al. 2007).

NLR helper pairs are relatively common. The *N. tabacum* NLR protein N requires an interaction with a second NLR protein named NRG1 to confer resistance to tobacco mosaic virus. Similarly, various ADR1 family proteins in *Arabidopsis* function as helper NLRs for resistance triggered by other NLRs (Peart et al. 2005; Bonardi et al. 2011). Many of these known ‘helper’ NLRs, including ADR1-L2 (Bonardi et al. 2011), contain P-loop mutations that inhibit the nucleotide binding necessary for canonical R protein activation (Bonardi et al. 2012). The *Arabidopsis* genome encodes many putative NLR proteins with degenerate P-loops, suggesting a widespread role for helper NLR proteins.

Elucidating the signaling mechanism used by RPS5 would shed light on whether the direct pore mechanism proposed for ZAR1 is a common signaling mechanism used by CC-NLRs, or whether the scaffolding mechanism used by mammalian NLRs is also present in plants. In either case, it could be that the signaling machinery used by RPS5 is a general mechanism used by other NLR proteins, like RPM1, that also do not re-localize upon activation (Gao et al. 2011).

The experiments above have shown that overexpression of PBS1^TuMV^ is capable of alleviating the trailing necrosis phenotype seen previously (Kim et al. 2016). This lends support to the correlation seen in Kim et al., 2016 between the expression levels of PBS1^TuMV^ and the corresponding severity of trailing necrosis. These observations indicate that the trailing necrosis phenotype observed by Kim et al. was due to suboptimal activation of RPS5, and that this can be overcome by increasing the amount of PBS1 available for cleavage, and hence increasing the amount of RPS5 activated above a threshold required to kill host cells prior to virus spread. It is noteworthy that overexpression of PBS1^TuMV^ or soybean PBS1^SMV^ did not cause observable changes in plant growth, which suggests that overexpression does not cause constitutive activation of defense responses, or other deleterious effects. Furthermore, it may be possible to express multiple PBS1 decoys in a single plant, further expanding the recognition specificity of RPS5.

The PBS1/RPS5 decoy system should be capable of conveying immunity to any biotrophic pathogen that requires a protease as part of its infection or replication machinery, providing that the protease accumulates inside the host cell and recognizes a cleavage sequence of seven, or fewer, amino acids. This has the potential to be broadly applicable as bacterial, viral and fungal pathogens, as well as oomycetes and nematodes, are known to employ proteases as part of their infection cycle (Alfano and Collmer 2004; Lim et al. 2011; Raffaele et al. 2010; Gardner et al. 2018).

This decoy system also has the potential to be transferred to crop plants without the need to transfer *At*RPS5, as many crop species are already resistant to *Pseudomonas syringae* through the recognition of AvrPphB, suggesting that they possess a functional analogue of RPS5 (Russell et al. 2015; Carter et al. 2019). Indeed, we have recently shown that soybean, barley and wheat all respond to AvrPphB protease activity (Carter et al. 2019; Helm et al. 2019), and this response in barley is mediated by a disease resistance protein designated Pbr1 (Carter et al. 2019).

*PBS1* is highly conserved in both dicots and monocots (Caldwell and Michelmore 2009; Swiderski and Innes 2001), including soybean, which contains three *PBS1* co-orthologs (Helm et al. 2019). Similar to the above work with Arabidopsis PBS1, insertion of an NIa cleavage site in the soybean PBS1 proteins enables their cleavage by the NIa protease from SMV (Helm et al. 2019). Significantly, co-expression of a decoy soybean PBS1 protein with the SMV NIa protease triggers cell death in soybean protoplasts (Helm et al. 2019). Our finding that transgenic overexpression of a soybean PBS1 decoy can confer complete resistance to SMV confirms that the PBS1 decoy approach can be deployed in crops using only endogenous genes. Thus, it should be feasible to engineer novel disease resistance traits in crops simply by making minor modifications to endogenous *PBS1* genes using genome editing approaches. Furthermore, in crops such as soybean and wheat, which have three or more *PBS1* co-orthologs, it should be possible to use genome editing to create multiple different PBS1 decoys that target different pathogens, or that target multiple proteases in a single pathogen to enhance durability.

## MATERIALS AND METHODS

### Plant growth

*Nicotiana benthamiana* and *Arabidopsis thaliana* plants were grown under a 12h light/12h dark cycle at 24°C in Pro-Mix PGX Biofungicide plug and germination mix supplemented with Osmocote 14-14-14 slow release fertilizer (Scotts).

Seeds of soybean [*Glycine max* (L.) Merr.] cultivars ‘Williams82’ and Williams82 *PBS1-1^SMV^* transgenic lines were sown in clay pots containing greenhouse soil (generated by composting plants and previously used potting mix, and then supplementing with small amounts of sand, perlite and vermiculite) supplemented with Osmocote slow-release fertilizer (14-14-14) and grown in a growth chamber under a 16 hr light/8 hr dark photoperiod at 23°C with average light intensities of 300 µEinsteins m^-2^ s^-1^ (at soil height).

### Generation of plant expression constructs

For the nuclear constructs, the RPS5 (AT1G12220) and PBS1^TuMV^ (based on PBS1 AT5G13160) open reading frames were amplified from plasmids described in Ade et al. 2007 and Kim et al. 2016 respectively, using PCR with primers that incorporated attB sites to enable Gateway™-based cloning (see Supplementary Table S1). In addition, the forward primers were designed to eliminate the native acylation motifs of PBS1 and RPS5. For NLS constructs, an NLS sequence (Gao et al. 2011) was incorporated into the forward primer. The 6K2-PBS1^TuMV^ construct was made through splicing overlap extension (SOE) PCR (Bryksin and Matsumura 2010). Separate PCR amplifications were performed to generate TuMV 6K2 and PBS1^TuMV^ fragments with the addition of attB sites at the 5’ end of the 6K2 sequence and the 3’ end of the PBS1 sequence for later cloning. The reverse primer of the 6K2 sequence included sequence that overlapped with the PBS1 sequence, which enabled joining of the two fragments via SOE PCR. The 6K2-RPS5 construct was produced by modifying an existing RPS5-5xMYC construct, as described in Ade et al. 2007, via In-Fusion cloning (Clontech) to fuse the 6K2 fragment (GenBank: D10927.1) to the N-terminus of RPS5. The resulting 6K2-RPS5-5xMYC construct (in plasmid pTA7001-DEST (Gu and Innes 2011)) was subjected to another round of PCR to isolate the 6K2-RPS5 fragment with the addition of attB sites for later cloning. The purified PBS1^TuMV^ and RPS5 fragments containing attB sites and NLS/KDEL/6K2 localization sequences were incorporated into pBSDONR-P1P4 using BP recombinase (Invitrogen). Constructs were recombined into the dexamethasone-inducible vector pBAV154 (Vinatzer et al. 2006) using multisite Gateway recombination cloning to add a superYFP tag for confocal microscopy and either a 3x HA tag, in the case of PBS1^TuMV^, or a 5x MYC tag, in the case of RPS5, for immunoblotting and cell death assays. Constructs were confirmed via sequencing before being transformed into *Agrobacterium tumefaciens* strain GV3101 (pMP90). The cytoplasmic RPS5 construct was previously detailed in Qi et al. 2012. The cytoplasmic PBS1^TuMV^ was made by mutating an existing cytoplasmic PBS1 construct in pBSDONR P1P4, as described in (Qi et al. 2014) to insert the TuMV NIa Protease consensus cleavage site. To generate the 35S:PBS1^TUMV^ construct, PBS1^TUMV^ in pBSDONR P1P4 and a C-terminal 3X HA tag were recombined into the destination vector pEG100 (Earley et al. 2006) using LR Clonase. The TuMV (6K2:GFP) construct was previously described by Wan et al. 2015. All primers used are listed in Supplementary Table S1, and all constructs generated are listed in Supplementary Table S2.

To generate the pWI-1000:E35S::*GmPBS1-1^SMV^-HA* construct, *GmPBS1-1^SMV^-HA* was PCR-amplified from the pBAV154:*GmPBS1-1^SMV^-HA* template (Helm et al., 2019) using primers designed to introduce *XbaI* restriction sites at each end. The resulting PCR products were gel-purified using the QIAquick gel extraction kit (Qiagen) and cloned into the *XbaI* site of pWI-1000 (M. Peterson; Wisconsin Crop Innovation Center). The resulting constructs were sequence-verified to check for proper sequence and reading frame.

### Generation of transgenic Arabidopsis lines

The 35S:6K2-PBS1^TuMV^-3xHA transgenic Arabidopsis lines were created by transferring the 6K2-PBS1^TuMV^ construct in pBSDONR P1P4 (Gu and Innes 2011) into pEG100 along with an 3xHA tag using multisite Gateway™ cloning. The resulting construct was confirmed via sequencing before being transformed into *Agrobacterium tumefaciens* strain GV3101 (pMP90). This strain was then used to prepare floral dips of *Arabidopsis rps5* and *pbs1* knockout lines, SALK lines 127201 and 062464 respectively (Clough and Bent 1998). Progeny were selected via BASTA resistance in T1 and T2 generations, and then homozygous T3 families identified. The same process was followed for the creation of 35S:PBS1^TuMV^-3xHA lines. The pbs1::PBS1^TuMV^-3xHA lines were described previously (Kim et al. 2016)

### Generation of transgenic soybean lines and infection with SMV

Transgenic Williams 82 soybean lines were generated by the Wisconsin Crop Innovation Center using an Agrobacterium-mediated transformation protocol (https://cropinnovation.cals.wisc.edu/services/soybean-transformation-2/). T1 seed from three independent T0 plants were obtained from the Wisconsin Crop Innovation Center. T1 plants were assessed for expression of a GUS (beta-glucuronidase) marker gene present on the T-DNA of the pWI-1000 vector using X-gluc staining of unifoliate leaves (Jefferson et al. 1987). T1 plants were selfed to obtain T2 seed. T2 plants were grown in a growth chamber at 22°C under a 16-hour day (300 µEinsteins m^-2^ s^-1^). Unifoliate leaves of 12-14 day old plants were rub-inoculated plasmid DNA encoding full-length SMV-NV::GFP (T1 plants), or SMV virions prepared from infected soybean plants (T2 plants), as described in Helm et al. 2019. In brief, virions were obtained from previously infected wild-type Williams 82 plants that were rub-inoculated with infectious pSMV-Nv:GFP plasmid DNA and maintained after infection for 4 weeks in a growth chamber. Heavily infected trifoliate leaves were harvested in 1X phosphate-buffered saline (137 mM NaCl, 2.7 mM KCl, 10 mM Na2HPO4, 1.8mM KH2PO4, pH 7.4), with cell debris cleared by centrifugation, and the supernatant containing virions stored at −80°C until use. Soybean plants were grown in a growth chamber until unifoliate leaves were fully expanded (approximately 12-14 days after planting). One of the two unifoliate leaves was wounded by rubbing carborundum on the abaxial side of the leaf. Using a clean cotton swab, virion suspension was rubbed on the abaxial side, and then plants were returned to the growth chamber for an additional three weeks. Third trifoliate leaves were harvested when completely expanded (approximately 21-days post inoculation), and photographed under white light and ultraviolet light. Tissue was flash frozen in liquid nitrogen and stored at −80°C.

### Transient protein expression

*Agrobacterium* cells were taken from LB plates and suspended in 10 mM MgCl_2_ with 100 μM acetosyringone (Sigma-Aldrich). After a two-hour incubation at room temperature, bacterial cultures were infiltrated into the leaves of three to four week old *N. benthamiana* with a needleless syringe. For microscopy and HR the final concentration of infiltrated bacteria, once mixed, was OD_600_ of 0.3 +/− 0.01, and for immunoblot analyses 0.15 +/− 0.01.

### TuMV Infection

*Agrobacterium* cells containing pCAMBIA carrying TuMV(6K2-GFP) (Wan et al. 2015) were taken from plates and suspended to an OD_600_ of 0.1 in 10 mM MgCl_2_ with 100 μM acetosyringone (Sigma-Aldrich). After a two-hour incubation at room temperature, bacterial cultures were infiltrated into three-week-old Arabidopsis leaves with a needleless syringe. Infection was allowed to progress for three weeks before being photographed under white and UV light.

### Fluorescence Microscopy

At 48 hours after infiltration, plants were sprayed with 50 µM dexamethasone, 0.02% Tween-20. Twenty-four hours post-transgene induction, leaves were imaged using a Leica SP5 scanning confocal light microscope and an HCX PL APO CS 63X 1.2 objective, pinhole 99.90 μM. Each construct was imaged at least three times, each from separate infiltration events.

### Immunoblots

For transient assays in *N. benthamiana,* 48 hours after infiltration, plants were sprayed with 50 µM dexamethasone. Six hours post-transgene induction, samples are taken for protein extraction. For assays in Arabidopsis, whole plat samples were taken immediately after TuMV symptoms were photographed. For samples in *N. benthamiana* leaves were pooled from two to three plants. Leaf tissue was ground in extraction buffer (150 mM NaCl, 50 mM Tris [pH 7.5], 0.2 % Nonidet P-40 [Sigma-Aldrich], 1% plant protease inhibitor cocktail [Sigma-Aldrich], 2 mM 2,2′-dipyridyl disulfide). Cell debris was pelleted at 8,000 x g, 4°C for 10 min, and the collected supernatants were separated on a 4-20% Tris-glycine Stain Free polyacrylamide gel (BioRad) at 185V for 1 hour in 1X Tris/Glycine/SDS running buffer. Total proteins were transferred to nitrocellulose membrane (GE Water and Process Technologies). Membranes were blocked overnight at 4°C in 5% milk. Proteins were subsequently detected with HRP-conjugated anti-HA antibody (rat monoclonal, Roche, catalog number 12013819001) or mouse monoclonal anti-GFP antibody (Novus Biologicals, Littleton, catalog number NB600-597). Membranes were washed 3 times for 15 mins in 1X Tris-buffered saline (TBS) with 0.1% Tween20 (TBST) and incubated with HRP-conjugated goat anti-mouse antibody (abcam, catalog number ab6789). All antibodies were used at a concentration on 1:5000. The nitrocellulose membranes were washed three times for 15 minutes in TBST and imaged using Supersignal® West Femto Maximum Sensitivity Substrates (Thermo Scientific) and X-ray film. The experiment was repeated three times.

For protein extraction from soybean, trifoliate leaves were ground to a powder under liquid nitrogen with mortar and pestle. Three times the volume of ice cold IP buffer (50 mM Tris-HCl, 150 mM NaCl, 10% glycerol, 1 mM diothioreitol, 1% NP-40, 0.1% Triton X-100, 1% Plant Protease Inhibitor cocktail, 1% DPDS, 1 mM EDTA) was added to powder and ground until homogenous. Leaf extracts were centrifuged 2x at 12,500xg at 4°C and supernatants free of plant tissue debris were collected. 500 µl of cleared supernatant was added to 10 µl anti-HA magnetic beads (MedChemExpress catalog number HY-K0201) and incubated on a rotator at 4°C for 3 hours. Immunoprecipitation samples were collected on a magnetic stand for 5 minutes and unbound protein lysate was removed and saved for SMV-CP analysis. Magnetic beads were washed 5 times in ice cold IP buffer. Protein was eluted in 40 µl of IP buffer and 10 µl of 5x SDS-PAGE buffer. Samples were boiled at 95°C for 5 minutes and returned to magnetic stand to separate sample from magnetic beads. A volume of 10 µl was loaded onto a 4-20% SDS-PAGE gel. 40 µl of unbound protein lysate was added to 10 µl of 5x SDS-PAGE loading buffer and boiled at 95°C for 5 minutes. Samples were briefly centrifuged and 10 µl of sample was loaded onto A 4-20% SDS-PAGE gel. Gels were run for approximately 1 hour at 160V and transferred to nitrocellulose membranes at 300 amps for 1 hour. Membranes with unbound protein lysates were stained with Ponceau for 3 minutes, rinsed with deionized water, and photographed for RUBISCO loading control.

SMV coat protein immunoblots were blocked overnight with 5% dry milk in 1x Tris-buffered saline (TBS; 50mM Tris-HCl, 150 mM NaCl, pH 7.5) solution containing 0.1% Tween 20 (TBST) and then anti-SMV coat protein antisera (rabbit polyclonal; a gift from Sue Tolin) added at a dilution of 1:20,000 and incubated at room temperature with gentle shaking for 1 hour. Membranes were washed three times with TBS-T, after which secondary goat anti-rabbit antibody conjugated to horse radish peroxidase (HRP) (Abcam, catalog number ab205718, Cambridge, MA) was added at a dilution of 1:5000 and incubated at room temperature for 1 hr, followed by washing three times with TBS-T. Anti-HA immunoblots were prepared in a similar fashion, but using an anti-HA-HRP conjugated antibody (rat monoclonal, Roche, catalog number 12013819001) at 1:5000, with incubation at room temperature for 1 hr.

Immunblots were developed using equal parts of ClarityTM Western ECL substrate peroxide solution and luminol/enhancer solution (BioRad) with incubation at room temperature for 5 minutes. Immunoblots were imaged using the chemiluminescent setting on the KwikQuant Imager (Kindle Biosciences, LLC).

### Cell death assays in *N. benthamiana*

At 48 hours after Agrobacterium infiltration into *N. benthamiana* leaves, plants were sprayed with 50 µM dexamethasone. Twenty-four hours later the infiltrated leaves were assessed for cell death and photographed under white light. Ten leaves over four to six plants were tested for every treatment in the experiment and the experiment was replicated three times.

### Cell death assays in Arabidopsis

*Pseudomonas syringae* strain DC3000 expressing either AvrPphB or an empty vector (Simonich and Innes 1995) were sub-cultured from plates into 10 µM MgCl_2_ and syringe infiltrated into leaves of four week old Arabidopsis. Twenty-four hours later the infiltrated leaves were assessed for cell death and photographed under white light.

### Aphid vector assays

Non-viruliferous aphid clones of a tobacco-adapted strain of *Myzus persicae* were reared under controlled conditions (23°C with a photoperiod of 12/12 h day/night) on tobacco (*Nicotiana tabacum*). Adult aphids were transferred to Col-0 leaves infected with TuMV (6k2-GFP) for a 10 min acquisition period. After virus acquisition, 2 aphids were transferred to a 20-day-old plantlet for each treatment for a 24 h inoculation period. After inoculation, the aphids were removed from the plants. Two weeks later, the number of infected plants was recorded for each line using a handheld UV light to visualize GFP, and tissue was collected for immunoblot confirmation. Seven to ten plants were inoculated per plant line.

For immunoblots of aphid-treated plants, 10 mg of leaf tissue was collected from each plant and pooled for each line. Leaf tissue was immediately frozen in liquid nitrogen and ground to a fine powder in a 1.5 mL tube using steel beads and a paint shaker. Lysis buffer (1 mL 0.5M Sodium citrate, 0.5 g SDS powder, 0.2 mL Beta-meracptoethanol, 1 mL 1.5M NaCl, 7.8 mL water, 1 tablet of EDTA-free Complete protease inhibitor cocktail) was added directly to the frozen tissue and mixed until homogeneous at room temperature (1 mg tissue:2 µL buffer). After mixing, samples were boiled for 10 minutes and cell debris pelleted in a microfuge at 15,000 RPM for 10 minutes at room temperature. Supernatants were mixed with loading buffer in a 1:1 ratio before separation at 70V for 15 minutes then 90V for 1.5 hours on a 10% Tris-glycine gel (Biorad) in 1X Tris/Glycine/SDS running buffer. Total proteins were transferred to a nitrocellulose membrane and blocked overnight at 4°C. Proteins were detected with anti-TuMV coat protein antibody at a concentration of 1:1500 and Goat anti-Rabbit IgG (H+L)-HRP conjugate (Miltenyi Biotec) at a concentration of 1:10,000, or using HRP-conjugated anti-HA antibody (Miltenyi Biotec) at a concentration of 1:2000. The nitrocellulose membranes were washed three times for 5, 15, and 10 minutes in TBST (1X TBS, 0.3% Tween) and imaged using a 1:4 ratio of Supersignal® West Femto Maximum Sensitivity Substrates and Supersignal® West Pico Plus (Thermo Scientific) and a ChemiDoc Imaging System (Biorad). Gels were stained with Coommassie blue (0.5g Coommassie R250, 200uL Methanol, 50uL Acetic Acid, 250uL Water) overnight at room temperature. Destaining solution (500mL water, 400mL methanol, 100mL Acetic Acid) was used for 2 hours. Coommassie blue stained gels were visualized using Gel Doc EZ Imager.

## ACKNOWLEDGEMENTS

We thank Dr. Jean-Francois Laliberte for providing the TuMV (6K2-GFP) construct; the Indiana University Light Microscopy Imaging Center for access to confocal microscopes; Leina Joseph, Indiana University, for technical assistance; Michael Peterson, Ray Collier, Mark Thompson, Amy Miyamoto, and Vai Sa Nee Lor at the Wisconsin Crop Innovation Center, Middleton, WI for generating the transgenic soybean lines; S. Tolin for generously providing antibody for the SMV coat protein; and the U.S. Department of Agriculture Soybean Germplasm Collection for soybean seed. This work was supported by a National Science Foundation Integrative Organismal Systems grant awarded to RWI (grant no. IOS-1551452). Mention of trade names or commercial products in this publication is solely for the purpose of providing specific information and does not imply recommendation or endorsement by the U.S. Department of Agriculture. USDA is an equal opportunity provider and employer.

