## Supplementary for "Optimizing the PBS1 Decoy System to Confer Resistance to Potyvirus Infection in Arabidopsis and Soybean"

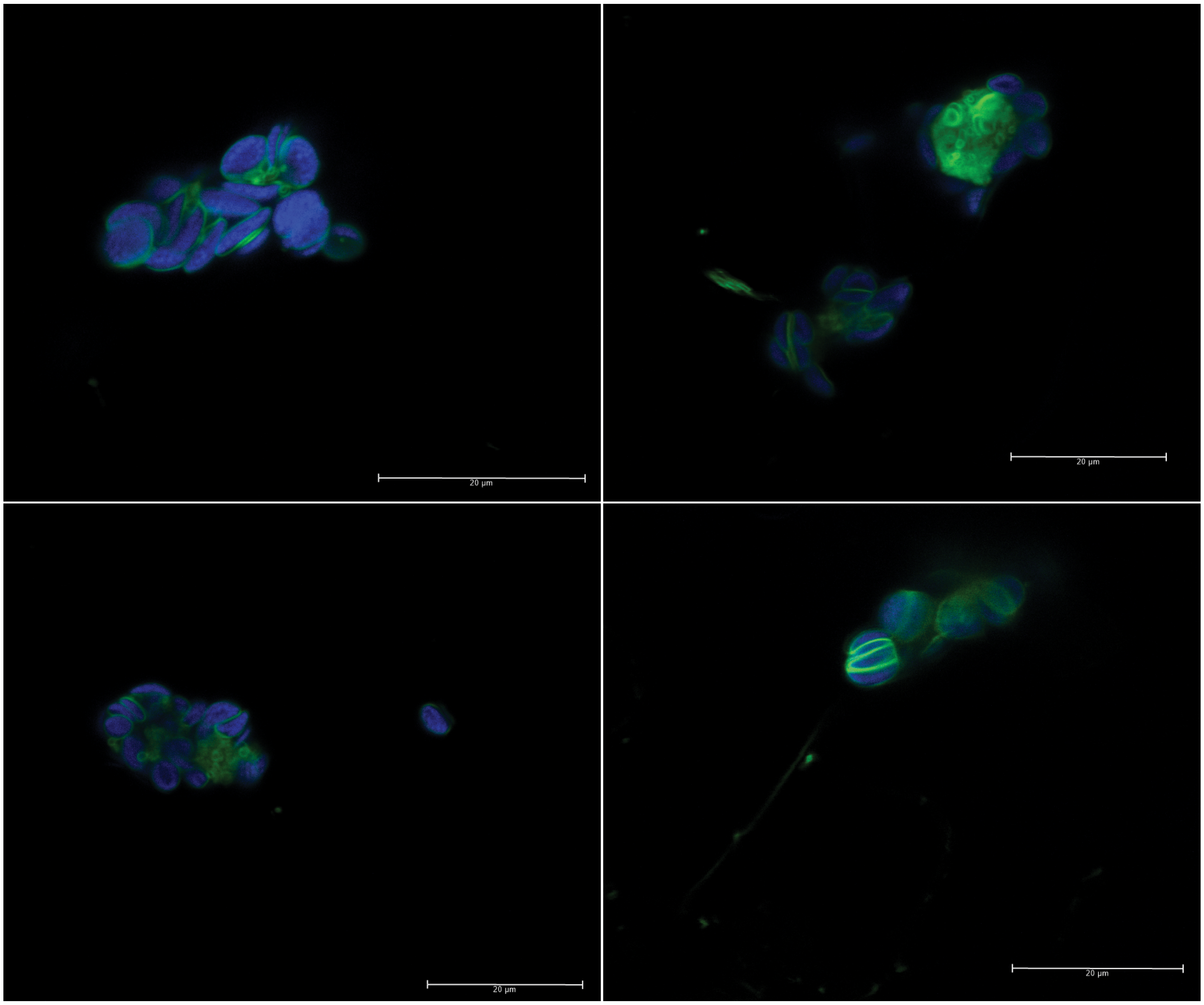


**Fig. S1.** Fluorescence images showing four representative chloroplast aggregations in *N. benthamiana* transiently expressing 6K2-PBS1$^{TuMV}$-YFP at 24 hours after the induction of gene expression. Chloroplast autofluorescence is shown in blue and 6K2-PBS1$^{TuMV}$-YFP is shown in green. The scale bar represents 20 µm.

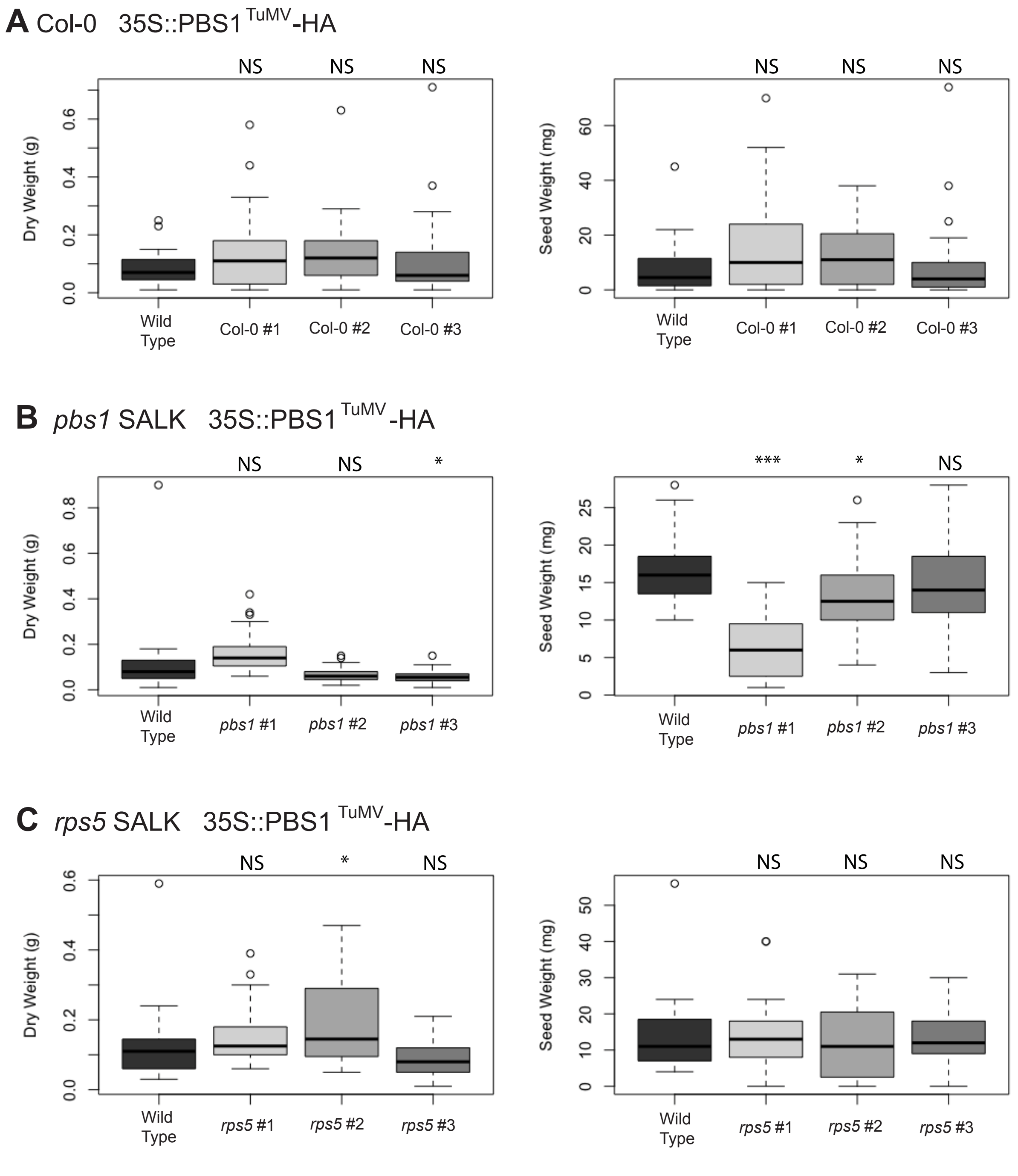


**Fig S2.** Box plots showing above ground dry weight and seed weight at senescence for each of the three 35S::PBS1$^{TuMV}$-HA transgenic lines in Col-0, *pbs1* SALK and *rps5* SALK backgrounds. Each set of transgenic lines is compared to WT Col-0 plants grown in the same tray. Analysis was done in R using a one-way ANOVA. * = $p<0.05$, ** = $p<0.01$, *** = $p<0.001$

**Supplementary Table S1.** Primers used in this study

| Primer Name | Sequence |
|---|---|
| AttB1 NLS-RPS5 FWD | GGGGACAAGTTTGTACAAAAAAGCAGGCTTAATGGGACCAAAGAAGAAACGGAAGGTCTTGCCATGTGATCAAGTAGTGAG |
| AttB4 NLS-RPS5 RVS | GGGGACAACTTTGTATAGAAAAGTTGGGTGTGTTTCTCTCCACCGCCA |
| AttB1 NLS-PBS1 FWD | GGGGACAAGTTTGTACAAAAAAGCAGGCTTAATGGGACCAAAGAAGAAACGGAAGGTC TCGAGTGATGACGAGAAGCTGA |
| AttB4 NLS-PBS1 RVS | GGGGACAACTTTGTATAGAAAAGTTGGGTGCCCGGTACTGTTGCTCTCTG |
| AttB4r RPS5-KDEL FWD | GGGGACAACTTTTCTATACAAAGTTGTAATGTTGCCATGTGATCAAGTAGTGAGT |
| AttB2 RPS5-KDEL RVS | GGGGACCACTTTGTACAAGAAAGCTGGGTTTTAGAGTTCATCCTTTGTTTCTCTCCACCGCCA |
| AttB4r PBS1-KDEL FWD | GGGGACAACTTTTCTATACAAAGTTGTAATGTCGAGTGATGACGAGAAGCTGA |
| AttB2 PBS1-KDEL RVS | GGGGACCACTTTGTACAAGAAAGCTGGGTTTTAGAGTTCATCCTTCCCGGTACTGTTGCTCTCTG |
| AttB1-6K2 FWD | GGGGACAAGTTTGTACAAAAAAGCAGGCTTAATGAGCACCAACGAAATGAGCAAG |
| 6K2-PBS1 RVS | TCAGCTTCTCGTCATCACTCGA TTCATGAGTTACGGGTTCAGA |
| 6K2-PBS1 FWD | TCTGAACCCGTAACTCATGAATCGAGTGATGACGAGAAGCTGA |
| AttB4-PBS1 RVS | GGGGACAACTTTGTATAGAAAAGTTGGGTGCCCGGTACTGTTGCTCTCTG |
| pTA7002-6K2 FWD | CGACTCTAGCTCGAGATGAGCACCAACGAAATGAGCAAG |
| 6K2-pTA7002/RPS5 RVS | TCCCATGTTCTCGAGTTCATGAGTTACGGGTTC |
| AttB1-6K2 FWD | GGGGACAAGTTTGTACAAAAAAGCAGGCTTAATGAGCACCAACGAAATGAGCAAG |
| AttB4-RPS5 RVS | GGGGACAACTTTGTATAGAAAAGTTGGGTGTGTTTCTCTCCACCGCCA |
| MUT-PBS1tumv FWD | CGGGAGGCTGTTCTCATCAGTCCACT |
| MUT-PBS1tumv RVS | AGTGGACTGATGAGAACAGCCTCCCG |
| GFP_GUS_FW | CAGCCATGCACACTGATA |
| GFP_GUS_RV | GATATCTACCCGCTTCGC |

**Supplementary Table S2.** Constructs generated for this study

| Constructs | | | | Use in Figure | | | | | | | | |
|---|---|---|---|---|---|---|---|---|---|---|---|---|
| **Location** | **Gene** | **Tag** | **Vector** | 1 | 2 | 3 | 4 | 5 | 6 | 7 | S1 | S2 |
| PM | PBS1tumv | YFP | pBAV154 | x | | | | | | | | |
| NLS- | PBS1tumv | YFP | pBAV154 | x | | | | | | | | |
| Cyt. | PBS1tumv | YFP | pBAV154 | x | | | | | | | | |
| 6K2 | PBS1tumv | YFP | pBAV154 | x | | | | | | | x | |
| PM | RPS5 | YFP | pTA7001 | x | | | | | | | | |
| NLS- | RPS5 | YFP | pBAV154 | x | | | | | | | | |
| -NLS | RPS5 | YFP | pTA7001 | x | | | | | | | | |
| Cyt. | RPS5 | YFP | pBAV154 | x | | | | | | | | |
| 6K2 | RPS5 | YFP | pBAV154 | x | | | | | | | | |
| PM | PBS1 | HA | pTA7001 | | | | | | x | | | |
| PM | PBS1tumv | HA | pBAV154 | | x | x | | | x | | | |
| PM | PBS1tumv | HA | pEG100 | | | | | x | x | x | | x |
| NLS- | PBS1tumv | HA | pBAV154 | | x | x | | | | | | |
| Cyt. | PBS1tumv | HA | pBAV154 | | x | x | | | | | | |
| 6K2 | PBS1tumv | HA | pBAV154 | | x | x | | | | | | |
| 6K2 | PBS1tumv | HA | pEG100 | | | | x | | | | | |
| PM | RPS5 | MYC | pBAV154 | | | x | | | | | | |
| NLS- | RPS5 | MYC | pBAV154 | | | x | | | | | | |
| -NLS | RPS5 | MYC | pBAV154 | | | x | | | | | | |
| Cyt. | RPS5 | MYC | pBAV154 | | | x | | | | | | |
| 6K2 | RPS5 | MYC | pBAV154 | | | x | | | | | | |
| PM | RPS5d266e | MYC | pBAV154 | | | x | | | | | | |
| NLS- | RPS5d266e | MYC | pBAV154 | | | x | | | | | | |
| -NLS | RPS5d266e | MYC | pBAV154 | | | x | | | | | | |
| Cyt. | RPS5d266e | MYC | pBAV154 | | | x | | | | | | |
| 6K2 | RPS5d266e | MYC | pBAV154 | | | x | | | | | | |
| | TuMV NIa | MYC | pBAV154 | | x | x | | | x | | | |
| | AvrPphB | MYC | pBAV154 | | | | | | x | | | |
| | 6K2 | GFP | pCAMBIA TuMV | | | | x | x | | x | | |
